# Impact of Parental Time-Restricted Feeding on Offspring Metabolic Phenotypic Traits

**DOI:** 10.1101/2024.06.04.597276

**Authors:** Yibo Fan, Xiangyuan Peng, Nishat I. Tabassum, Xiangru Cheng, Sharmelee Selvaraji, Vivian Tran, Tayla A. Gibson Hughes, Buddhila Wickramasinghe, Abdulsatar Jamal, Quynh Nhu Dinh, Mathias Gelderblom, Grant R. Drummond, Christopher G. Sobey, Jim Penman, Terrance G. Johns, Raghu Vemuganti, Jayantha Gunaratne, Mark P. Mattson, Dong-Gyu Jo, Maria Jelinic, Thiruma V. Arumugam

## Abstract

A substantial body of research elucidates the mechanisms and health advantages associated with intermittent fasting (IF). However, the impact of parental IF on offspring remains unclear. Through an investigation involving four IF and *ad libitum* combinations of parental mating groups, we demonstrate that parental IF (daily time-restricted feeding) influences offspring’s metabolic health indicators in male and female offspring in distinct ways. We found that when both parents are on IF their offspring exhibit protection against the adverse effects of a high-fat, high-sugar, and high-salt diet in a sex-specific manner. This study underscores the potential significance of parental lifestyle modifications involving dietary restriction for the metabolic status of their children and their risk for obesity and diabetes.

## Background

Over the past nine decades, a multitude of studies exploring the effects of dietary restriction, encompassing caloric restriction (CR) and IF, have consistently revealed their ability to enhance metabolic health and confer resistance to a range of chronic diseases. CR and IF not only offer protective effects against age-related ailments but also exhibit positive impacts on cognition and cellular resilience against various environmental challenges.

Recent research has shown that dietary restriction induces epigenetic modifications and associated changes in gene expression in animals^**1-4**^. Our investigations have further elucidated this link, demonstrating the impact of IF on the epigenome^**5,6**^. We found that IF modulates histone trimethylation, thereby orchestrating a myriad of transcriptomic changes that drive robust metabolic switching processes^**5**^. Moreover, a portion of the epigenomic and transcriptomic changes induced by IF persist even after refeeding, suggesting the maintenance of memory for such epigenetic changes^**5**^. Similarly, global DNA methylation patterns in IF animals distinctly deviate in CG nucleotide-rich islands (CGIs)^**6**^. These findings suggest the potential for heritable epigenetic changes accompanying the phenotypic response to IF in parental animals.

Despite the well-documented health benefits of IF in both animals and humans^**7**^, the precise nature and extent of IF-induced epigenetic changes, and the prospect of intergenerational inheritance of such changes are unknown. It is therefore of interest to explore whether IF represents an environmental trigger capable of modulating phenotypic profiles not only in parents but also in their descendants. In this study, we investigated the intergenerational effects of IF in one or both parents on the metabolic phenotypes of their offspring.

## Results

Parental male and female C57BL/6 mice were all initially fed a normal diet on a caloric basis until the start of the dietary intervention. At 6 weeks of age, mice were randomly assigned to either *ad libitum* (AL) feeding or a daily 16-hour fasting (IF) schedule. The study design, including the timing of experimental interventions and blood and tissue collections, is outlined in **Supplementary Fig. 1**. Parental mice on AL feeding exhibited significantly greater body weight (**Supplementary Fig. 2a and b)** and body weight changes (**Fig. 1a and b)** than those on the IF, regardless of sex, during the 16-week dietary intervention period. To evaluate the impact of IF on energy metabolism in parental animals, we quantified blood glucose, hemoglobin A1c (HbA1c), ketone, cholesterol, triglyceride, and the inflammation marker C-reactive protein (CRP) levels at baseline and at 8 and 16 weeks of dietary intervention (**Fig. 1c-j; Supplementary Fig. 3**). Male and female mice on IF exhibited significantly lower glucose levels (**Fig. 1c and d**) and HbA1c levels (**Supplementary Fig. 3a and b**), and significantly higher ketone levels (**Fig. 1e and f**) compared to the AL control mice. Furthermore, cholesterol levels were significantly reduced in IF mice from both sexes at week 16 (**Fig. 1g and h**). Triglyceride levels were similar in IF and AL mice (**Supplementary Fig. 3c and d**). CRP levels were significantly reduced by IF in female mice at 16 weeks (**Fig. 1i and j**).

**Figure 1:**
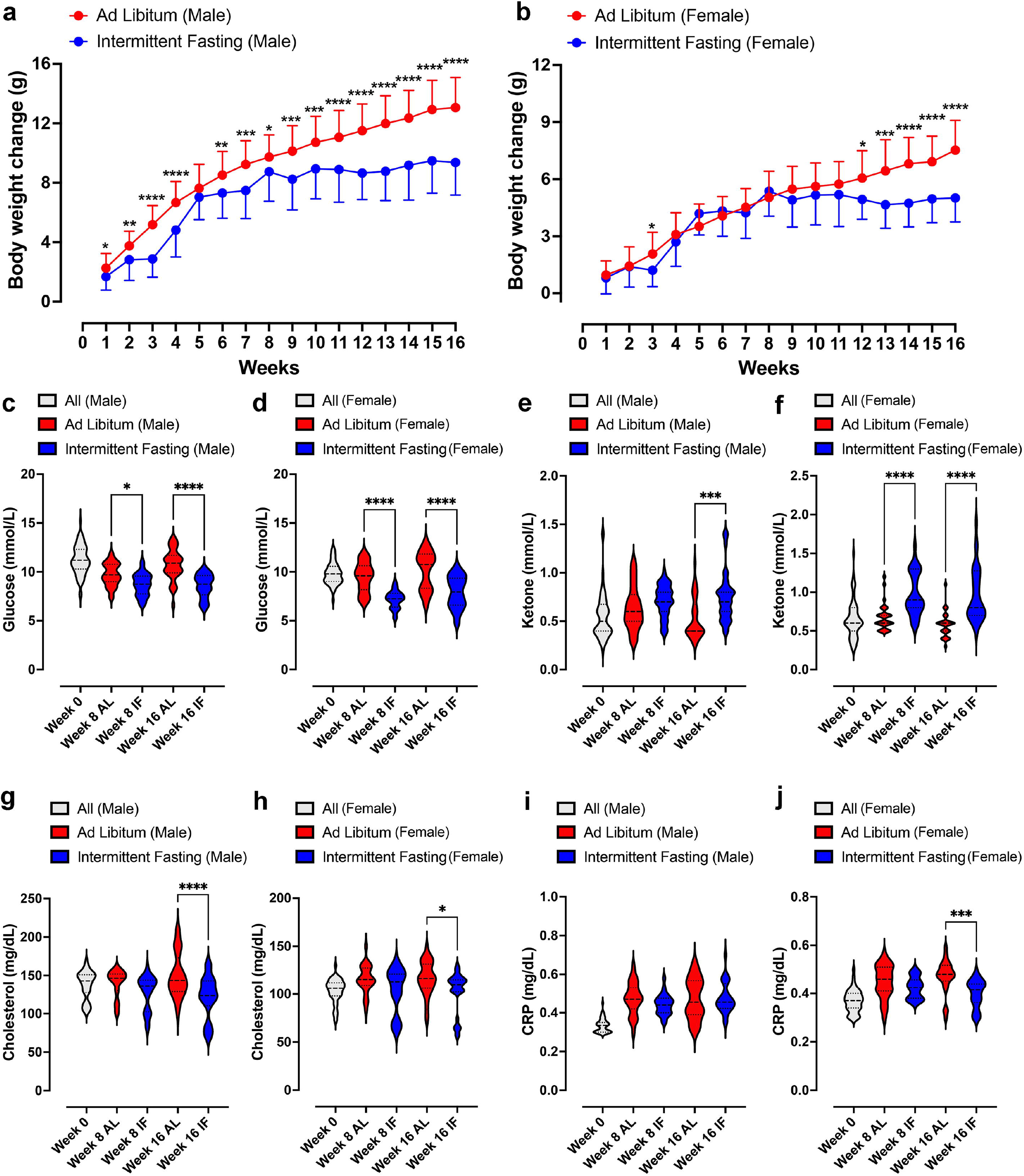
Impact of Intermittent Fasting on Parental Animals. (a) and (b) show the change in body weight in male and female animals, respectively, following daily 16-hour intermittent fasting (IF) over a 16-week period. Data are presented as means ± standard deviation. Statistical analysis was conducted using two-way repeated-measures ANOVA, followed by post-hoc Šídák’s multiple comparisons test. Significance levels are denoted as *P < 0.05, **P < 0.01, ***P < 0.001, ****P < 0.0001 compared to IF animals. Each group consisted of n = 32 animals. (c) and (d) display violin plots representing blood glucose levels in male and female animals, respectively. (e) and (f) depict blood ketone levels in male and female animals, while (g) and (h) show cholesterol levels. (I and j) illustrates C-reactive protein levels in male and female parental animals. Statistical analysis for these parameters was conducted using one-way ANOVA, followed by post-hoc Šídák’s multiple comparisons test. Significance levels are denoted as *P < 0.05, ***P < 0.001, ****P < 0.0001 compared to *ad libitum* (AL) animals. Each group consisted of n = 32 animals.

After 16 weeks of dietary intervention, parental male and female mice from both groups were paired for mating. During the mating period, all mice had *ad libitum* access to food and water. Upon confirmation of pregnancy, male mice were separated from the cages, and female mice had unrestricted access to food and water throughout pregnancy and a subsequent 4-week period.

When they were 4 weeks old the offspring from all four parental groups, namely F_1_(F_0_ AL Male X F_0_ AL Female); F_1_(F_0_ AL Male X F_0_ IF Female); F_1_(F_0_ IF Male X F_0_ AL Female); and F_1_(F_0_ IF Male X F_0_ IF Female), were separated from their mothers and assigned to either the usual normal diet or a high-fat, high-sugar, and high-salt (HFSS) diet (see **Supplementary Fig. 1**). The offspring were then maintained on either a normal or HFSS diet for an additional 16 weeks. Throughout this period, they underwent evaluations for physiological changes and metabolic phenotypes to investigate potential effects of maternal and/or paternal IF. Notably, male offspring on the normal diet from parents that had both been on IF exhibited a significant reduction in body weight from week 10 onwards compared to offspring from parents that had both been on AL (**Supplementary Fig. 4a and b**). Female offspring from parents that had both been on IF exhibited a trend towards lower average weight under both normal and HFSS diets (**Supplementary Fig. 4e and g**).

We next measured blood glucose and insulin levels in all groups of offspring after an intraperitoneal glucose tolerance challenge (**Figs. 2 and 3**). This glucose tolerance test assesses how effectively the body clears a glucose load. For males on the normal diet, those from an AL female and IF male crossing exhibited a significantly greater glucose area under the curve (AUC) compared to offspring from AL female/AL male crossings. When on the normal diet males from IF female/IF male crossings exhibited a lower AUC compared to males from AL female/AL male crossings (**Fig. 2a and b**). However, no significant differences were observed in blood glucose AUC levels among female offspring on the normal diet between any of the four different combinations of parental groups (**Fig. 2c and d**). For male offspring on the HFSS diet, there were no significant differences in glucose AUC among between any of the four parental diet groups (**Fig. 2e and f**). However, female offspring on the HFSS diet, from IF male/IF female crossings exhibited significantly lower glucose AUC levels compared to offspring from each of the other three groups (**Fig. 2g and h**).

**Figure 2:**
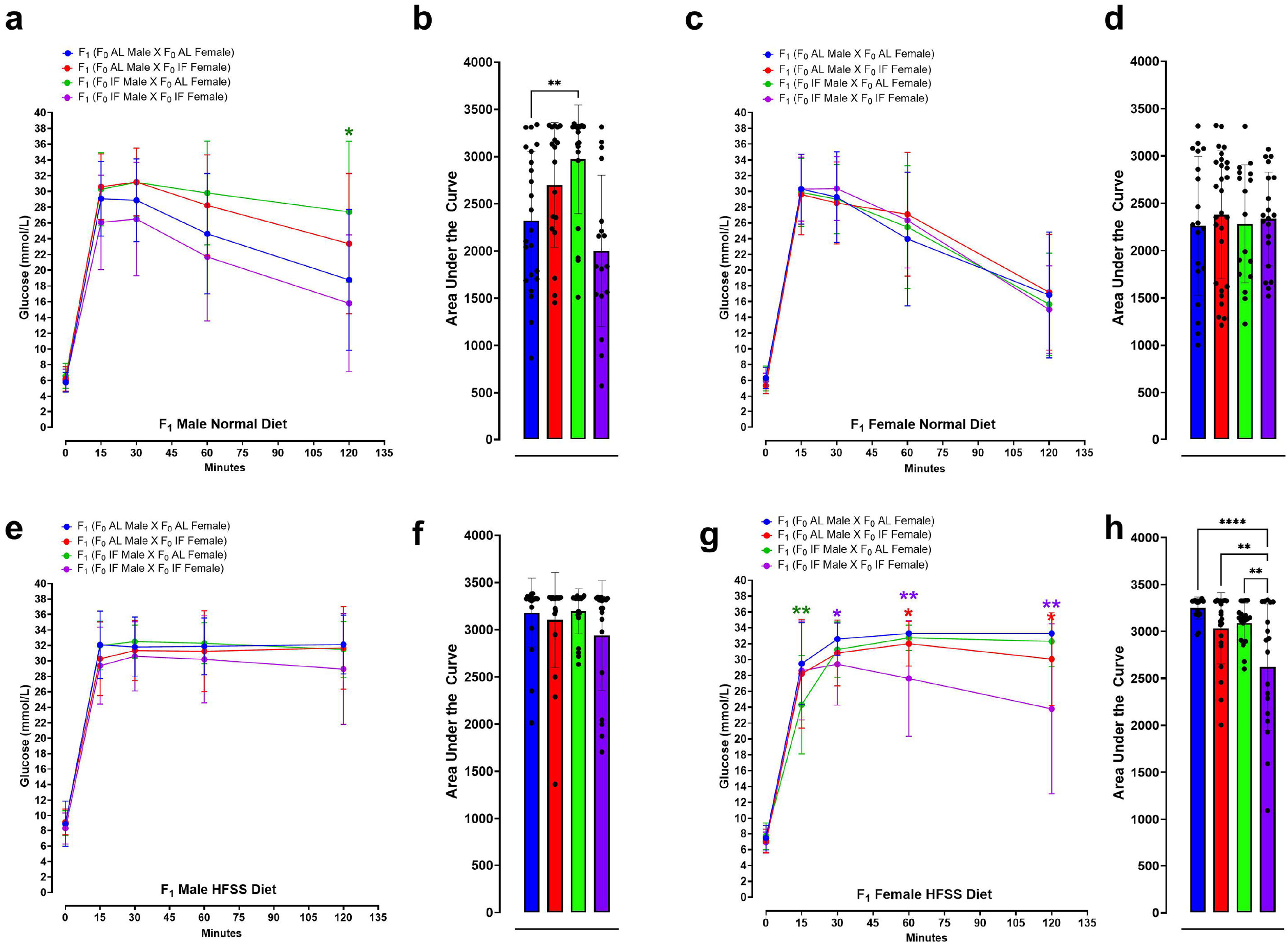
Glucose Tolerance Test Analysis in Offspring. Blood glucose concentrations during glucose tolerance test in normal diet male (a and b) and normal diet female (c and d) F_1_ mice at 0, 15, 30, 60 and 120 minutes post intraperitoneal glucose injection. Statistical analysis was conducted using two-way repeated-measure ANOVA, followed by post-hoc uncorrected Fisher’s LSD comparisons test. *P < 0.05 versus normal diet F_1_ (F_0_ AL male X F_0_ AL Female) animals. (b and d) Glucose area under the curve (AUC) for normal diet male (b) and normal diet female (d) F_1_ mice. Statistical analysis was performed using one-way ANOVA, followed by Tukey’s multiple comparisons test. **P < 0.01, versus F_1_ (F_0_ AL male X F_0_ AL Female) animals. Each group consisted of n = 17-28 animals. Similar analyses were conducted for F_1_ mice fed a high-fat, high-sugar and high-salt (HFSS) diet. Blood glucose concentrations during glucose tolerance test in HFSS diet male (e and f) and HFSS diet female (g and h) F_1_ mice at 0, 15, 30, 60 and 120 minutes after intraperitoneal glucose injection. Statistical significance was assessed using two-way repeated-measure ANOVA, followed by post-hoc uncorrected Fisher’s LSD comparisons test. *P < 0.05, **P < 0.01 indicate significance compared to *P < 0.05, **P < 0.01 versus F_1_ (F_0_ AL male X F_0_ AL Female) animals. Glucose AUC for HFSS diet male (f) and HFSS diet female (h) F_1_ mice. One-way ANOVA, followed by Tukey’s multiple comparisons test, **P < 0.01, ****P < 0.0001 versus HFSS F_1_ (F_0_ AL male X F_0_ AL Female) animals. Each group consisted of n = 18-22 animals.

**Figure 3:**
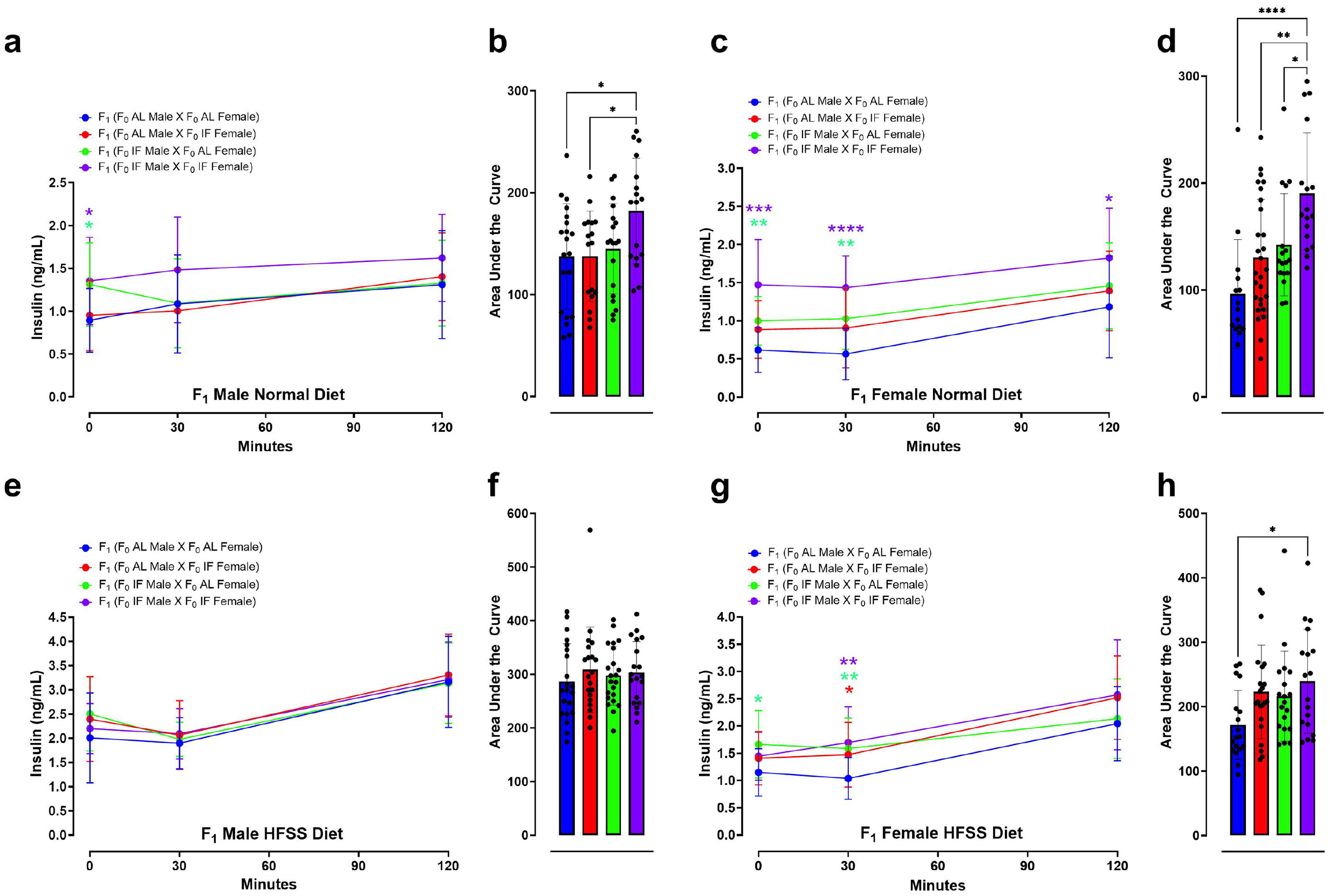
Insulin Response in Offspring Following Glucose Tolerance Test. Blood insulin levels during the glucose tolerance test in male (a and b) and female (c and d) F_1_ mice fed a normal diet, measured at 0, 30, and 120 minutes post intraperitoneal glucose injection. Statistical analysis was conducted using two-way repeated-measure ANOVA, followed by post-hoc uncorrected Fisher’s LSD comparisons test. *P < 0.05, **P < 0.01, ***P < 0.001, ****P < 0.0001 denote significance compared to normal diet F_1_ (F_0_ AL male X F_0_ AL Female) animals. Insulin AUC for male (b) and female (d) F_1_ mice fed a normal diet was calculated. Statistical analysis was performed using one-way ANOVA, followed by Tukey’s multiple comparisons test. *P < 0.05, **P < 0.01, ****P < 0.0001 indicate significance compared to F_1_ (F_0_ AL male X F_0_ AL Female) animals. Each group consisted of n = 15-28 animals. Similar analyses were conducted for F1 mice fed a high-fat, high-sugar, high-salt (HFSS) diet. Blood insulin concentrations during the glucose tolerance test in HFSS diet male (e and f) and female (g and h) F_1_ mice were measured at 0, 30, and 120 minutes post injection. Statistical significance was assessed using two-way repeated-measure ANOVA, followed by post-hoc uncorrected Fisher’s LSD comparisons test. *P < 0.05, **P < 0.01 denote significance compared to F_1_ (F_0_ AL male X F_0_ AL Female) animals. Insulin AUC for HFSS diet male (f) and female (h) F1 mice was determined, with statistical analysis conducted using one-way ANOVA, followed by Tukey’s multiple comparisons test. *P < 0.05 denotes significance compared to HFSS F_1_ (F_0_ AL male X F_0_ AL Female) animals. Each group consisted of n = 18-22 animals.

We measured plasma insulin levels at baseline (0 minutes; before peritoneal glucose injection) and at 30- and 120-minutes thereafter. For mice on the normal diet, insulin AUC analysis revealed significantly elevated insulin levels in male offspring of male IF and female IF crossings compared to males from AL/AL or male AL/female IF crossings (**Fig. 3a and b**). For offspring females on the normal diet those from IF male/IF female crossings exhibited elevated insulin AUC levels compared to female offspring from each of the other three groups (**Fig. 3c and d**). For offspring males on the HFSS diet, there were no differences in insulin AUC levels among the four groups (**Fig. 3e and f**). For offspring females on the HFSS diet those from IF male/IF female crossings exhibited an elevated insulin AUC level compared to those from AL male/AL female crossings (**Fig. 3g and h**). For offspring on the normal diet, the insulin levels obtained just before the intraperitoneal glucose injection were significantly higher in male and female offspring from IF male/IF female crossings compared to offspring from AL male/AL female crossings (**Fig. 3a and c**).

Plasma glucose, ketone, cholesterol, triglyceride, and CRP levels were measured immediately following weaning (Week 0), and at 12 and 16 weeks of dietary intervention preceding the glucose tolerance test (**Supplementary Figures 5, 6 and 7**). As expected, mice on the HFSS diet exhibited elevated glucose levels compared to those on the normal diet (**Supplementary Figure 5**). Offspring males and females from IF male/IF female crossings had lower plasma glucose levels compared to those from AL male/AL female crossings. Ketone levels were overall similar in mice on normal and HFSS diets. Offspring females from IF male/IF female crossings exhibited significantly higher ketone levels compared to offspring females from AL male/AL female crossings (**Supplementary Fig. 5**). Mice on the HFSS diet exhibited elevated cholesterol levels compared to those on the normal diet (**Supplementary Figure 6**). Offspring males on the normal diet from IF male/IF female crossings had lower basal plasma cholesterol levels compared to those from AL male/AL female crossings. Similarly, basal plasma triglycerides levels were lower in offspring males on the normal diet from IF male/IF female crossings and IF male/AL female crossings compared to offspring females from AL male/AL female crossings (**Supplementary Figure 6**). Mice on the HFSS diet exhibited elevated CRP levels compared to those on the normal diet (**Supplementary Figure 7**).

## Discussion

Previous studies have shown that parental nutritional status can significantly impact offspring through epigenetic inheritance mechanisms^**8-12**^. While some research has explored the phenotypic outcomes in offspring resulting from parental obesogenic diets^**13,14**^, the possible effects of parental IF on offspring metabolic traits are unknown. The present study begins to fill this knowledge gap. We found that compared to offspring from parents fed AL, those born to parents that had both been on IF exhibit differences in markers of energy metabolism and that vary depending upon the sex of the offspring. Male and female parents on IF exhibited lower plasma glucose and HbA1c levels and elevated ketone levels compared to those fed AL. On the normal diet offspring males from IF/IF parents had significantly lower body weights compared to offspring males from AL/AL parents, whereas offspring female body weight was unaffected by parental IF. Offspring females (but not offspring males) from IF/IF parents exhibited improved glucose tolerance when on a HFSS diet but not when on the normal diet.

Our findings suggest that parental fasting leads to enhanced insulin production both at basal levels (minute 0 in both female and male offspring animals under normal diet) and following glucose injection, indicating that these offspring inherit epigenetic mechanisms that may be responsible for the observed increase in insulin secretion. Previous studies have established that glucose metabolism is significantly improved with dietary restriction^**15-17**^. Notably, while fasting insulin concentrations were reduced in db/db mice, they were maintained in db/db mice under CR^**18**^. db/db mice are used to model diabetes type II and obesity that are homozygous for the diabetes spontaneous mutation (*Lepr*^*db*^) demonstrate morbid obesity.

In conclusion, our study represents a novel exploration into how parental IF may confer metabolic benefits to offspring, likely mediated by epigenetic mechanisms inherited from both parents. Notably, female offspring of IF parents, but not males, retained these metabolic advantages even when exposed to an HFSS diet. To delve deeper into this inheritance and its sex-specific manifestation, we have initiated comprehensive genomic investigations. While our results offer valuable insights, it’s crucial to acknowledge the inherent metabolic disparities between species, raising caution in directly translating our observations to humans.

## Methods

### Animals and IF procedures

All *in vivo* experimental procedures were approved by La Trobe University (Ethics approval number: AEC21047) Animal Care and Use Committees and conducted in accordance with the guidelines outlined in the Australian Code for the Care and Use of Animals for Scientific Purposes (8th edition) and the NIH Guide for the Care and Use of Laboratory Animals. Every effort was made to minimize suffering and reduce the number of animals used. All sections of the manuscript were performed in accordance with ARRIVE guidelines. C57BL/6 male and female mice were purchased at 4 weeks of age from ARC, Australia, and housed in the animal facility at La Trobe University. The animals were subjected to a 12-hour light:12-hour dark cycle (07:00-19:00) and provided with normal diet comprising 20% total crude protein, 5% crude fat, 6% crude fiber, 0.5% added salt, 0.8% calcium and 0.45% phosphorus (Ridley). Water was available *ad libitum* to all dietary groups. At 6 weeks of age, male and female mice were randomly assigned to different dietary intervention groups: Intermittent Fasting for 16 hours (IF16; n=64) and *ad libitum* feeding as a control (AL; n=64). Mice in the IF16 group underwent daily fasting for 16 hours for a duration of 4 months (16:00-08:00), while the AL group had continuous access to food pellets. Body weight was regularly monitored, and blood glucose and ketone levels were measured using the FreeStyle Optium Neo system with FreeStyle Optium blood glucose and ketone test strips (Abbott) 6 hours after fasting. Additionally, cholesterol (Roche Cobas b 101 Lipid), triglycerides (Roche Cobas b 101 Lipid), HbA1c (Roche Cobas b 101 HbA1c) and CRP (Roche Cobas b 101 CRP) levels were assessed at different time points from all animals after 6 hours fasting (at the onset of dietary interventions, as well as at 8 and 16 weeks post-intervention). After 4 months of dietary intervention, animals were assigned to mating groups. After the mating process and subsequent weaning of offspring were completed, all parental animals were euthanized by carbon dioxide (CO2) inhalation between 7 a.m. and noon. Subsequently, all mice were perfused with cold PBS, and tissue samples were collected and snap-frozen in liquid nitrogen and stored at -80 °C until further use.

### Mating and offspring production

After 4 months of dietary intervention, male and female mice were randomly divided into four groups: AL fathers, IF fathers, AL mothers, and IF mothers. The mating groups were assigned as follows: AL-father X AL-mother; AL-father X IF-mother; AL-mother X IF-father; IF-father X IF-mother. The offspring were obtained and divided into male and female groups upon weaning, with 36-50 animals per group (8 groups in total). Furthermore, both female and male offspring were further divided into normal diet groups and high-fat food and high-salt, and high-sugar drinking water (HFSS; High-fat: 20.90% protein, 23.50% total fat, 5.40% crude fiber, 5.40% AD fiber; SF04-001, Specialty Feeds, Australia; High-salt (0.9% Sodium Chloride, Baxter); High-sugar (4.5% glucose and 5.5% fructose) groups, with 17-28 animals per group (16 groups in total). Please refer to **Figure 1** for details on grouping. All offspring were placed on their assigned dietary regimen for a period of 16 weeks. Body weight was weekly measured. Blood glucose and ketone levels were assessed in the morning using the FreeStyle Optium Neo system with FreeStyle Optium blood glucose and ketone test strips before IF animals received food. Additionally, cholesterol, triglycerides, HbA1c and CRP levels were measured: at the start of different dietary interventions, as well as at 8 and 16 weeks after the interventions.

### Glucose tolerance tests and insulin measurement

Glucose tolerance tests were performed following an overnight fasting period of 16±2 hours, with basal blood glucose and plasma insulin samples (t = 0) measured. Mice were administered 2 grams of glucose per kilogram of body weight via intraperitoneal injection using a 20% glucose solution (BioUltra). Blood glucose levels were monitored at 15, 30, 60, and 120 minutes post-glucose challenge. Additional blood samples were obtained in 0.8 ml microtube with LH Lithium Heparin Separator (MiniCollect) for plasma insulin analysis at 30 and 120 minutes post-glucose injection. To prepare plasma samples, blood samples were centrifuged at 10, 000 RPM for 10 minutes at 4°C (Eppendorf), followed by transfer of the plasma to tubes. These tubes were then snap-frozen in liquid nitrogen and stored at −80°C for subsequent insulin measurements. Insulin levels were quantified using Mouse Ultrasensitive Insulin ELISA (ALPCO), with each sample assayed in duplicate following the manufacturer’s instructions.

### Statistical analysis

All analyses were performed using GraphPad Prism version 10.0. For Figure 1 and Supplementary Figure 1 (F_0_ generation), as well as Supplementary Figure 3 (F_1_ generation), body weight and changes in body weight data were presented as means ± standard deviation (s.d.). For all comparisons involving multiple time points in Figure 1 (F_0_ generation) and Supplementary Figure 2 (F_0_ generation), a two-way repeated-measurements analysis of variance (ANOVA), followed by post-hoc Šídák’s multiple comparisons test, was conducted to determine P values compared to control groups. For other parameters depicted in Figure 1 (F_0_ generation), data were displayed using box plots and violins, and a one-way ANOVA, followed by post-hoc Šídák’s multiple comparisons tests, was used to determine P values compared to respective time-specific control groups. For Figures 2 and 3 (F_1_ generation), in the analysis of the glucose and insulin levels following the tolerance test, two methods were employed. Firstly, the XY graph depicts individual replicates with means connected, and a two-way repeated-measurements ANOVA, followed by post-hoc uncorrected Fisher’s LSD comparisons test, was utilized to determine P values compared to respective time-specific control groups. Secondly, the Area Under the Curve analysis was performed using one-way ANOVA, followed by Tukey’s multiple comparisons test, to determine P values. Data were presented as means ± standard deviation (s.d.). For Supplementary Figures 4-6 (F_1_ generation), data were presented as means ± standard error of the mean (s.e.m.). A two-way repeated-measurements ANOVA was conducted, followed by post-hoc uncorrected Tukey’s multiple comparisons test, to determine P values.

## Data availability

All source data will be made available upon request.

## Contributions

T.V.A. and Y.F. designed the study, analyzed data, and co-wrote the manuscript. X.P., N.I.T., X.C., V.T., T.A.G.H., B.W., A.J., and Q.N.D. contributed to data collection. S.S., M.G., G.R.D., C.G.S., J.P., T.G.J., R.V., and J.G. contributed to intellectual discussions on the study design or provided experimental support throughout the study. M.M. and D.G.J. contributed to the original intellectual design of the study. M.J. developed the HFSS method applied in this study and provided intellectual input into the study design. All authors participated in writing the manuscript. T.V.A. supervised the study. All authors reviewed and approved the final manuscript.

## Ethics declarations

All *in vivo* experimental procedures were approved by La Trobe University (Ethics approval number: AEC21047) Animal Care and Use Committees and performed according to the guidelines set forth by the Australian Code for the Care and Use of Animals for Scientific Purposes (8th edition) and confirmed NIH Guide for the Care and Use of Laboratory Animals.

---

**Supplementary Figure 1: Experimental Design for Parental Animals and Offspring.** Parental animals were divided into two groups: one received *ad libitum* (AL) access to food, while the other underwent daily 16-hour intermittent fasting (IF16) for 4 months before mating. All parental mice were raised under AL conditions for 2 months before being randomly assigned to either continue AL feeding or undergo IF16. Offspring were separated from their mothers at 4 weeks of age and further divided into male and female groups. These groups were then subdivided into those receiving a normal diet or a high-fat, high-sugar (HFSS) diet, and were maintained under these dietary conditions for 16 weeks.

**Supplementary Figure 2: Impact of Intermittent Fasting on Body Weight of Parental Animals.** The figure illustrates the body weight trajectories of male (a) and female (b) parental animals following daily 16-hour intermittent fasting (IF) over a 16-week period. Data are presented as means ± standard deviation. Statistical analysis was conducted using two-way repeated-measure ANOVA, followed by post-hoc Šídák’s multiple comparisons test. Significance levels are indicated as *P < 0.05, **P < 0.01, ***P < 0.001, ****P < 0.0001 compared to IF animals. Each group consisted of n = 32 animals.

**Supplementary Figure 3: Impact of Intermittent Fasting on HbA1c and Triglyceride Levels of Parental Animals.** Violin plots depict blood HbA1c levels in male (a) and female (b) parental animals, as well as triglyceride levels in male (c) and female (d) parental animals. Statistical analysis was conducted using one-way ANOVA, followed by post-hoc Šídák’s multiple comparisons test. Significance levels are denoted as *P < 0.05, ****P < 0.0001 compared to *ad libitum* (AL) animals. Each group consisted of n = 32 animals.

**Supplementary Figure 4: Body Weight of Offspring Animals.** The figure presents the body weight trajectories of male offspring (a and c) and female offspring (e and g) under normal and high-fat, high-sugar, high-salt (HFSS) diets over 16-week periods. Data are displayed as means ± standard deviation. Statistical analysis was conducted using two-way repeated-measurement ANOVA, followed by post-hoc Šídák’s multiple comparisons test. Significance levels are indicated as *P < 0.05 compared to F_1_ (F_0_ AL male X F_0_ AL Female) animals. Each group comprised n = 17-28 animals. Violin plots illustrate the body weight distributions of offspring normal diet male (b), HFSS diet male (d), normal diet female (f) and HFSS diet female (h) animals at 16 weeks after weaning. Statistical analysis was performed using one-way ANOVA, followed by post-hoc Šídák’s multiple comparisons test. Significance levels are denoted as *P < 0.05 compared to F_1_ (F_0_ AL male X F_0_ AL Female) animals. Each group consisted of n = 17-28 animals.

**Supplementary Figure 5: Blood Glucose and Ketone Levels of Offspring Animals.** Bar graphs illustrate the blood glucose and ketone levels at week 0 (immediately after weaning), week 12, and week 16 in male offspring under normal diet (a and b), high-fat, high-sugar, high-salt (HFSS) diet (c and d), as well as female offspring under normal diet (e and f) and HFSS diet (g and h). Statistical analysis was conducted using one-way ANOVA, followed by post-hoc Tukey’s multiple comparisons test. Significance levels are denoted as *P < 0.05, **P < 0.01. Each group comprised n = 17-28 animals.

**Supplementary Figure 6: Blood Cholesterol and Triglyceride Levels of Offspring Animals.** Bar graphs depict the blood cholesterol and triglyceride levels at week 0 (immediately after weaning), week 12, and week 16 in male offspring under normal diet (a and b), high-fat, high-sugar, high-salt (HFSS) diet (c and d), as well as female offspring under normal diet (e and f) and HFSS diet (g and h). Statistical analysis was performed using one-way ANOVA, followed by post-hoc Tukey’s multiple comparisons test. Significance levels are indicated as *P < 0.05, **P < 0.01, ****P<0.0001. Each group consisted of n = 17-28 animals.

**Supplementary Figure 7: Blood CRP Levels of Offspring Animals.** Bar graphs illustrate the blood CRP levels at week 0 (immediately after weaning), week 8, and week 16 in male offspring under normal diet (a), high-fat, high-sugar, high-salt (HFSS) diet (b), as well as female offspring under normal diet (c) and HFSS diet (d). Statistical analysis was conducted using one-way ANOVA, followed by post-hoc Tukey’s multiple comparisons test. Significance levels are denoted as *P < 0.05. Each group comprised n = 18-28 animals.

