## Supplementary figures and images for "Impact of Parental Time-Restricted Feeding on Offspring Metabolic Phenotypic Traits"

### Supplementary Figure 1

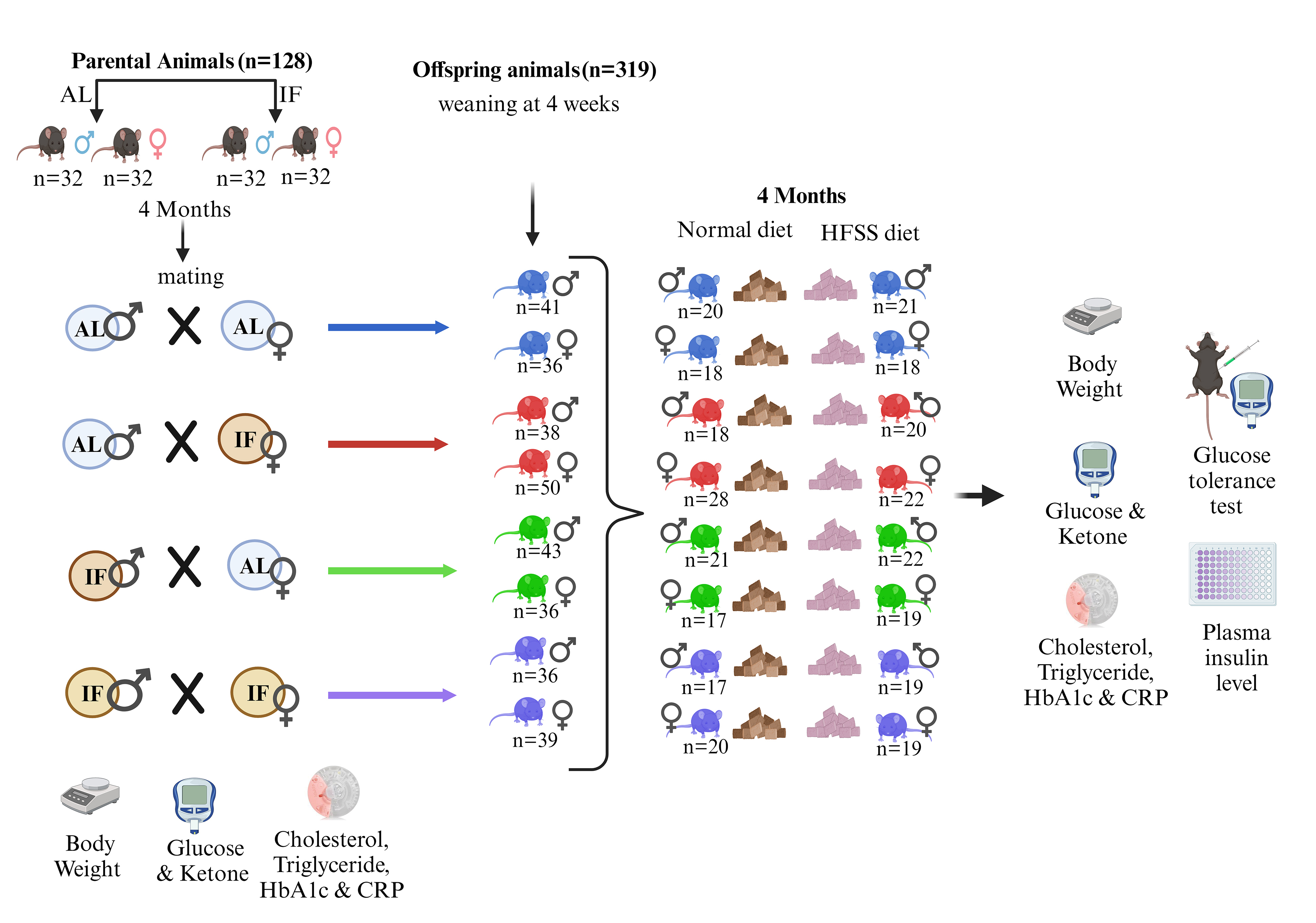

### Supplementary Figure 2

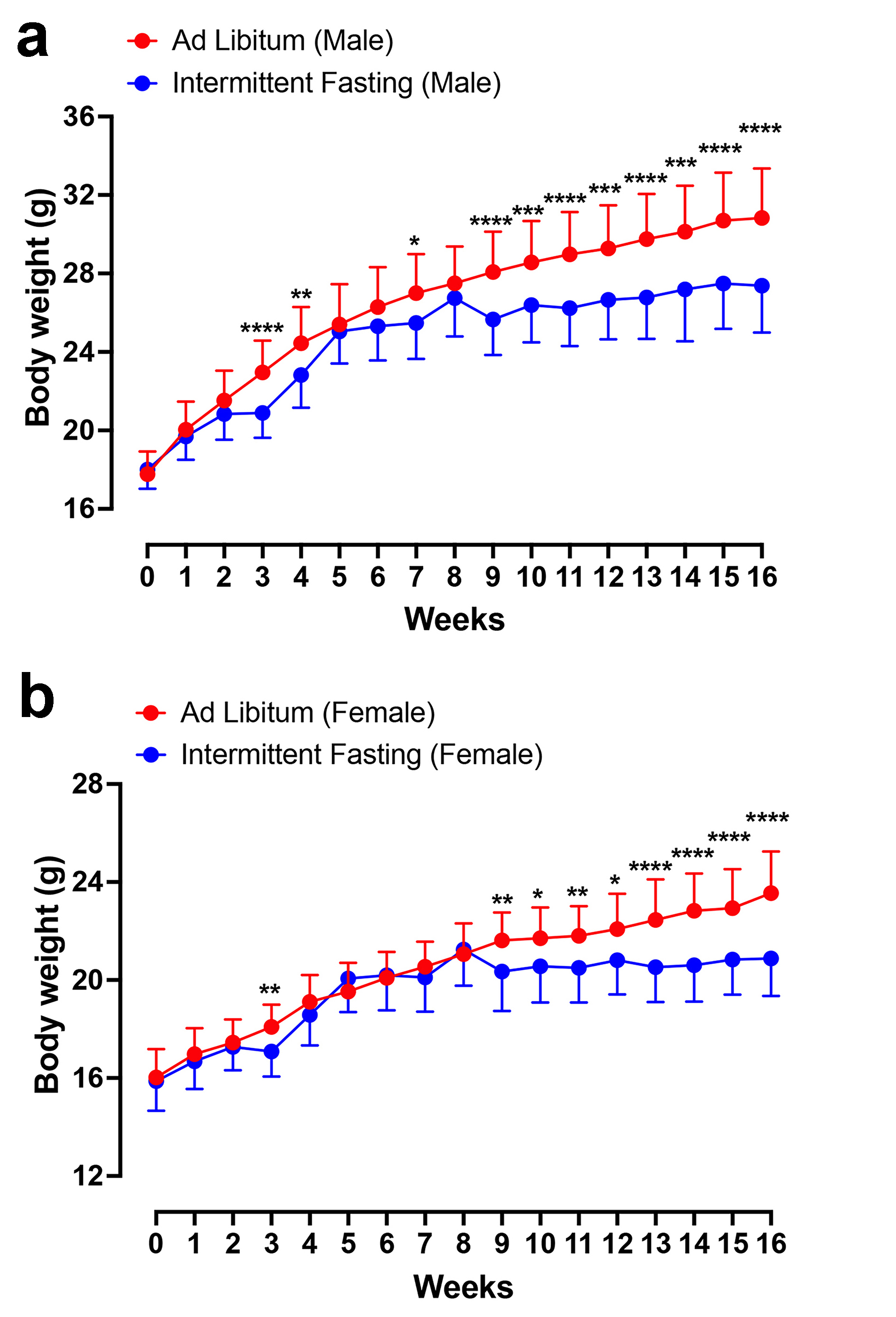

### Supplementary Figure 3

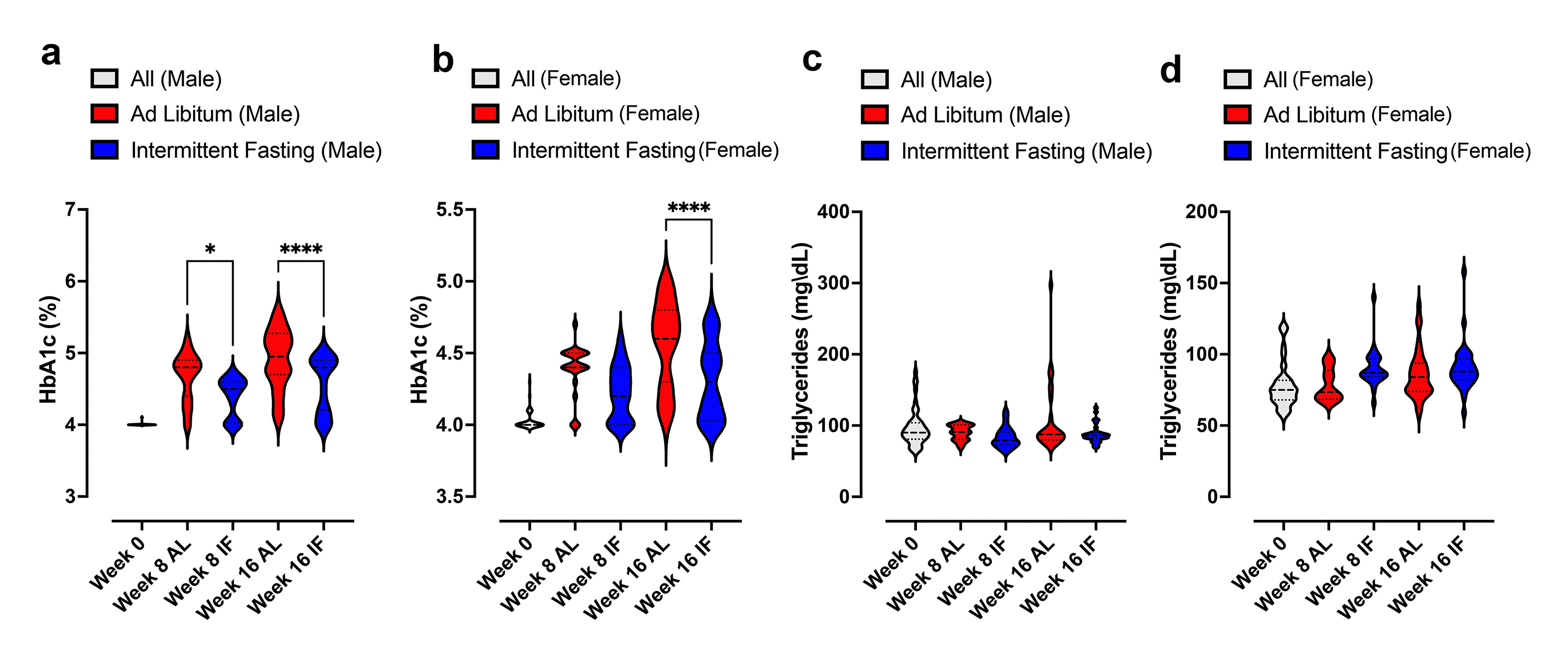

### Supplementary Figure 4

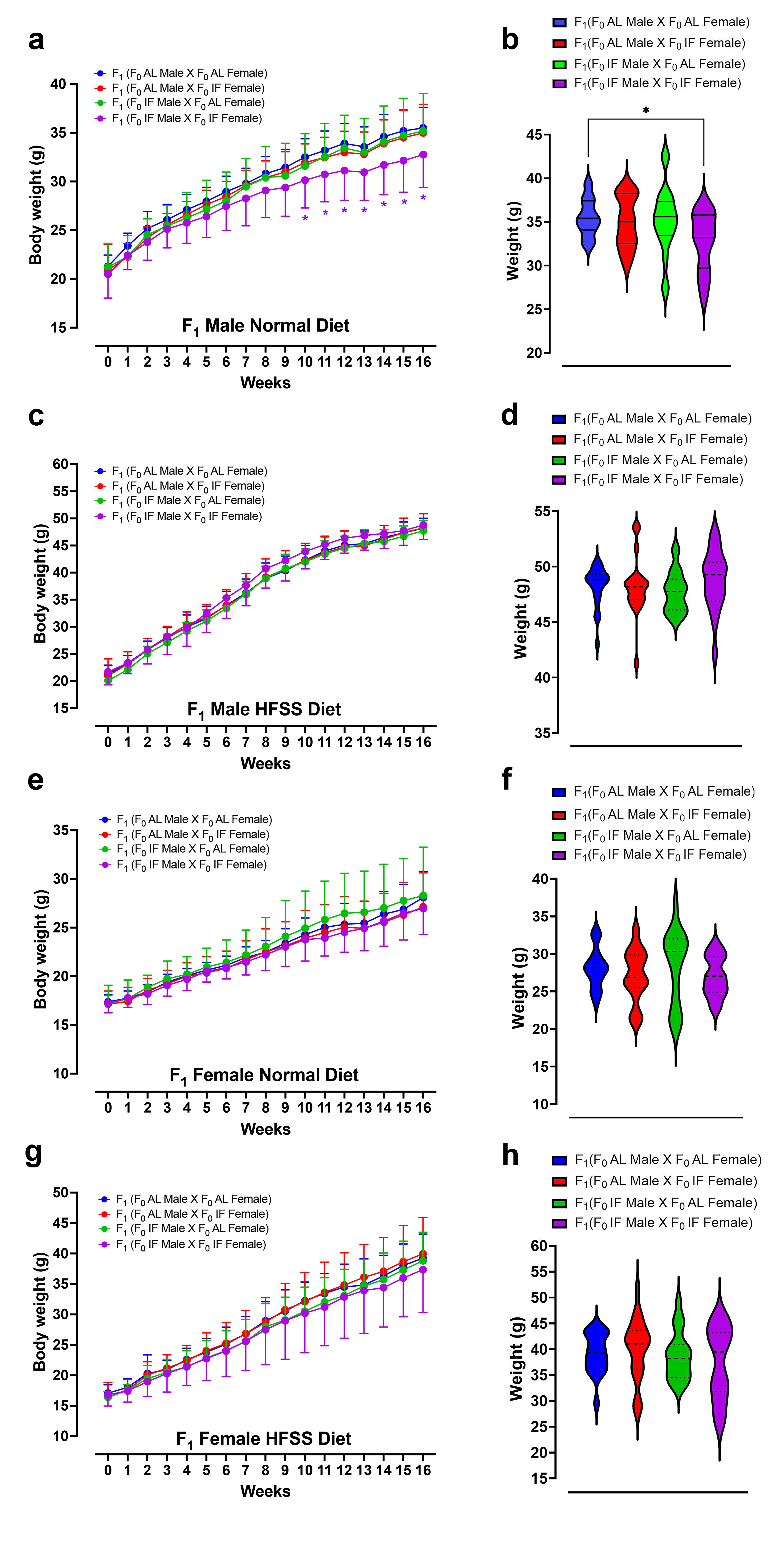

### Supplementary Figure 5

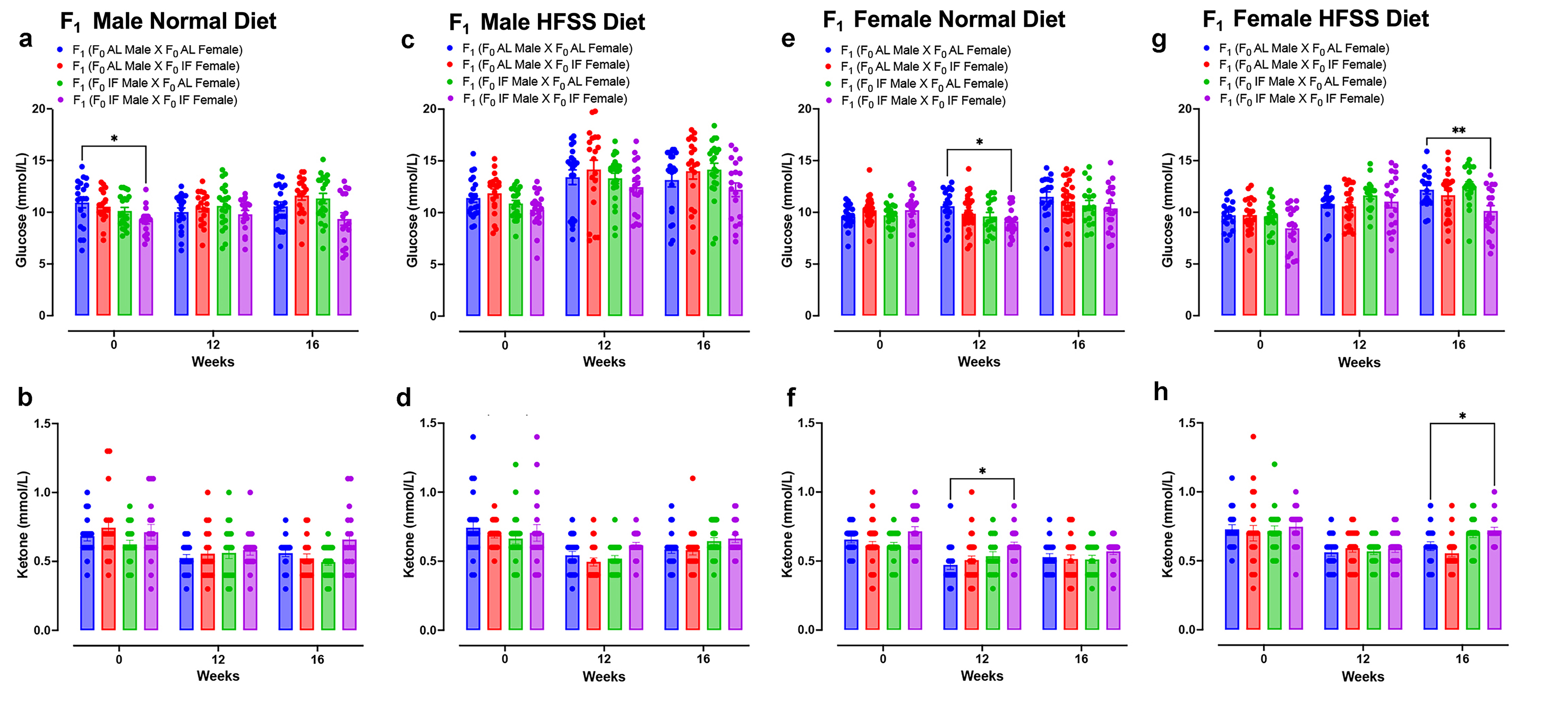

### Supplementary Figure 6

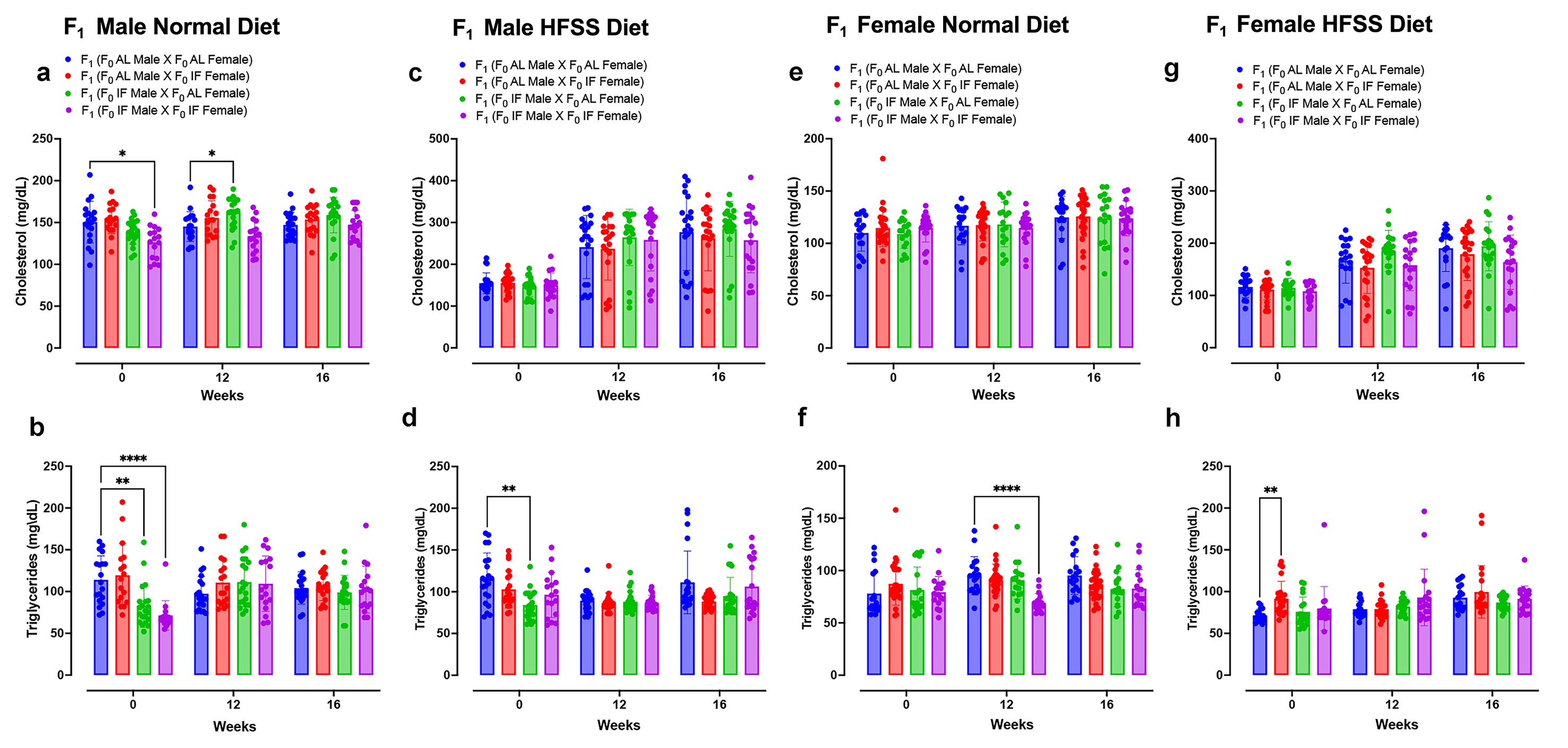

### Supplementary Figure 7

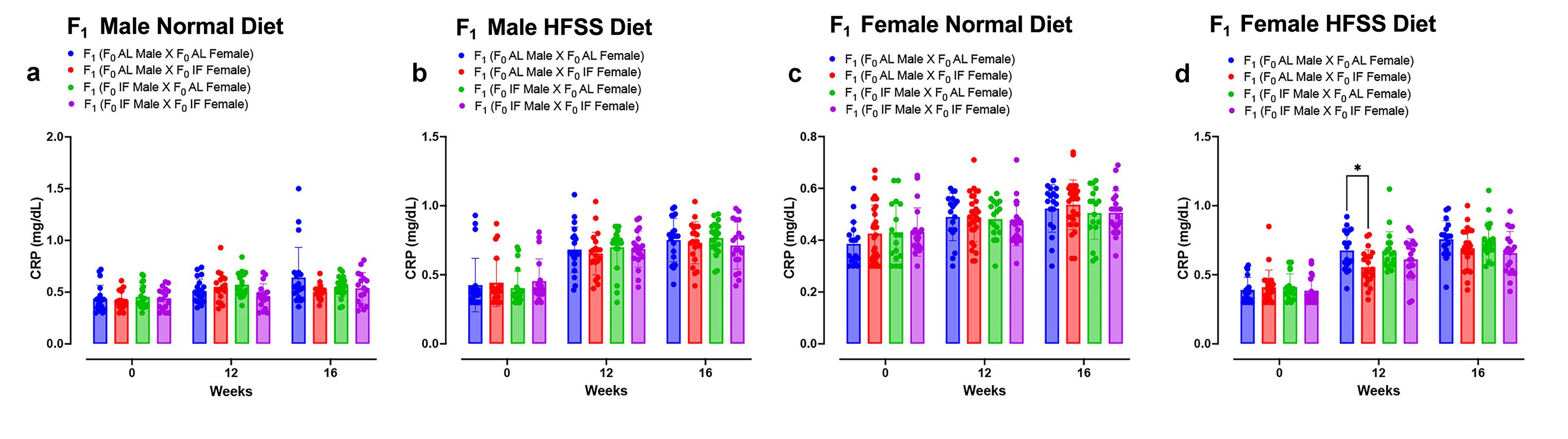
